# The CURE for Cultivating Fastidious Microbes

**DOI:** 10.1101/167130

**Authors:** Arundhati Bakshi, Austen T. Webber, Lorelei E. Patrick, E. William Wischusen, J. Cameron Thrash

## Abstract

Course-based Undergraduate Research Experiences (CUREs) expand the scientific educational benefits of research to large groups of students in a course setting. As part of an ongoing effort to integrate CUREs into first-year biology labs, we developed a microbiology CURE (mCURE) that uses a modified dilution-to-extinction high throughput culturing protocol for isolating abundant yet fastidious aquatic bacterioplankton during one semester. Students learn common molecular biology techniques like nucleic acid extraction, PCR, and molecular characterization; read and evaluate scientific literature; and receive training in scientific communication through written and oral exercises that incorporate social media elements. In the first three semesters, the mCUREs achieved similar cultivability success as implementation of the protocol in a standard laboratory setting. Our modular framework facilitates customization of the curriculum for use in multiple settings and we provide classroom exercises, assignments, assessment tools, and examples of student output to assist with implementation.

## INTRODUCTION

Undergraduate research experiences in STEM increase student retention in science majors; increase the proportion of students that go on to professional or graduate school; as well as improve critical thinking skills, data interpretation skills, content knowledge, and attitudes toward science (1-5). Typical undergraduate research experiences are limited to relatively few students due to research lab size and funding, making these positions competitive, highly selective, and typically dominated by upper-level students (4, 5). Course-based undergraduate research experiences (CUREs), in which students experience research as part of a course, can reach students early in their degree program and accommodate large numbers of students, thus increasing the diversity of students participating in research (4, 5). Despite these benefits, the time necessary to plan CURE projects and create assignments and rubrics can restrict their use (6). Fortunately, an increasing number of publications have shared CURE implementation strategies for a variety of settings (3, 7-9). We recently outlined a flexible, modular CURE framework, including rubrics and course materials, that has facilitated conducting a variety of different research projects in first-year biology laboratory courses at Louisiana State University (LSU) (10). Using this framework, we have developed the microbiology CURE (mCURE) described herein that focuses on the cultivation of bacterioplankton from aquatic systems (**Fig. 1**).

**Figure 1.**
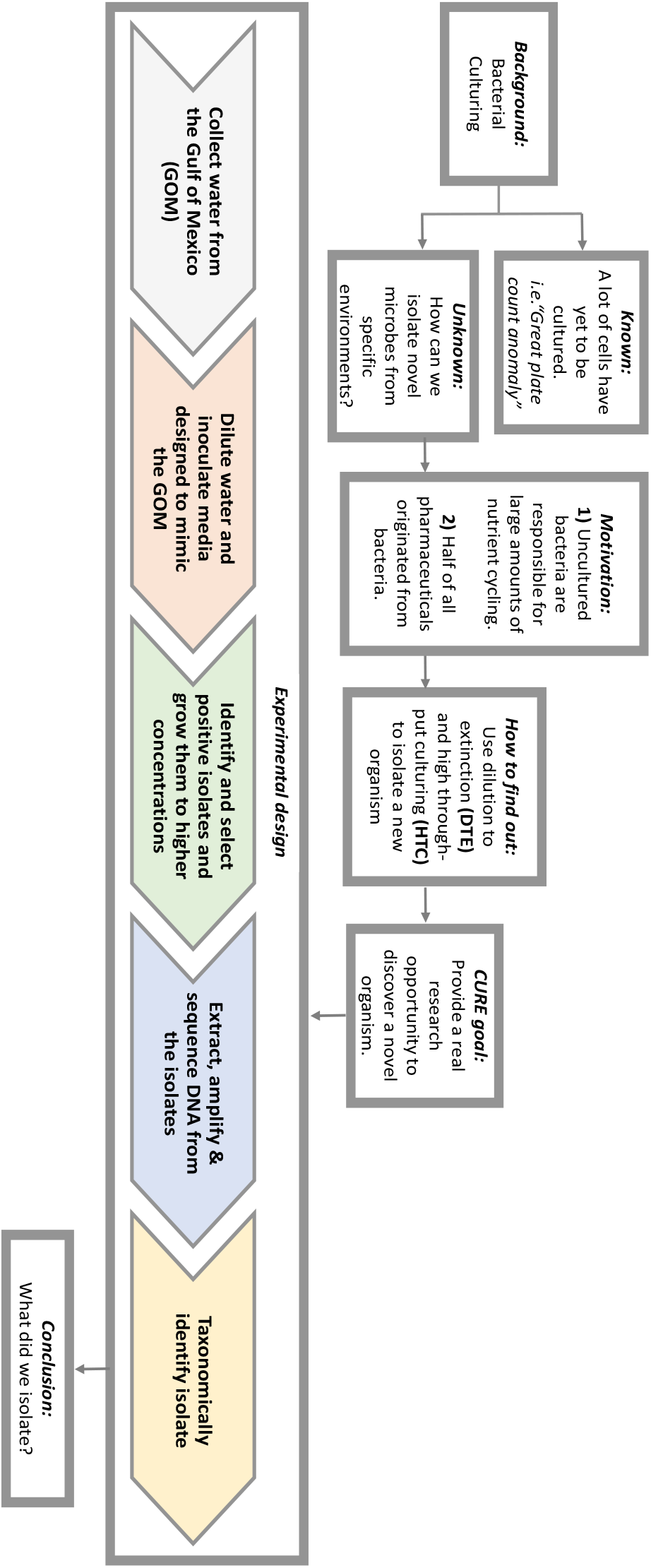
Flowchart of the mCURE background and experimental design. Using this flowchart, students are guided through the scientific process to gain an understanding of the relevance and importance of the project. Various segments of the course are color-coded (grey, orange, green, blue and yellow) corresponding to Table 1 where the week-by-week activities for each of these segments are described. This flowchart may be modified as needed to suit alternative projects using a similar protocol.

Bacterioplankton occupy marine and freshwater environments at cell concentrations typically between 10^5^-10^7^ cells mL^-1^, however traditional agar plate methods usually only cultivate 0.1-1% of the organisms present in a given sample (11), hampering our ability to understand the functions of a large majority of microorganisms. An improved high-throughput cultivation (HTC) method combines serial dilution of samples with sterilized natural water and/or artificial seawater media (12-14). Many abundant taxa in aquatic systems have been successfully cultured using this approach, for example SAR11 *Alphaproteobacteria* (15-18), SUP05 *Gammaproteobacteria* (19), SAR116 *Alphaproteobacteria* (12, 20), and members of the so-called “Oligotrophic Marine *Gammaproteobacteria*” (21). Artificial media facilitates more general application and modification (e.g., in salinity, carbon and nitrogen sources, etc.) to accommodate different environments, as well as the adaptation of the protocol to teaching laboratories. In the following mCURE, students execute a modified version of the HTC protocol utilized by the Thrash Laboratory at LSU (14, 22). The possibility of isolating new organisms provides a charismatic entrance into biological research where students experience a genuine excitement of discovery combined with their laboratory and communication training.

### Intended audience

This course teaches basic laboratory skills and molecular biology methods, such as DNA extraction and PCR, in the context of advanced microbial cultivation approaches and introduces students to identification of microorganisms with molecular techniques. The curriculum also includes exercises in reading and understanding primary literature and communicating science to different audiences. The course is intended for undergraduates at the first- or second-year level who are pursuing majors such as Biology and Microbiology.

### Learning time

We designed the mCURE for a semester timeline with a single three-hour laboratory section meeting once a week for a minimum of 13 weeks. The project is divided into four major segments (color-coded in both **Figure 1** and **Table 1**). In weeks #2-4 (***orange***), students attempt to establish an initial culture of marine bacterioplankton using serial dilutions with the HTC protocol (22). Transfer of the initial cultures to larger flasks for further growth occurs during weeks #5-6 (***green***). During weeks #7-9 (***blue***), students extract DNA from the cultures and amplify the 16S rRNA gene with PCR. Amplified products are then sequenced for subsequent taxonomic identification of the microbes in week #10 (***yellow***). The remaining weeks (#11-13) are spent discussing poster construction and administering the final assessments. Note that the entire workflow does not require 13 weeks, but we have built in flexibility to allow for repeating one or more elements in case of failure.

**Table 1.**
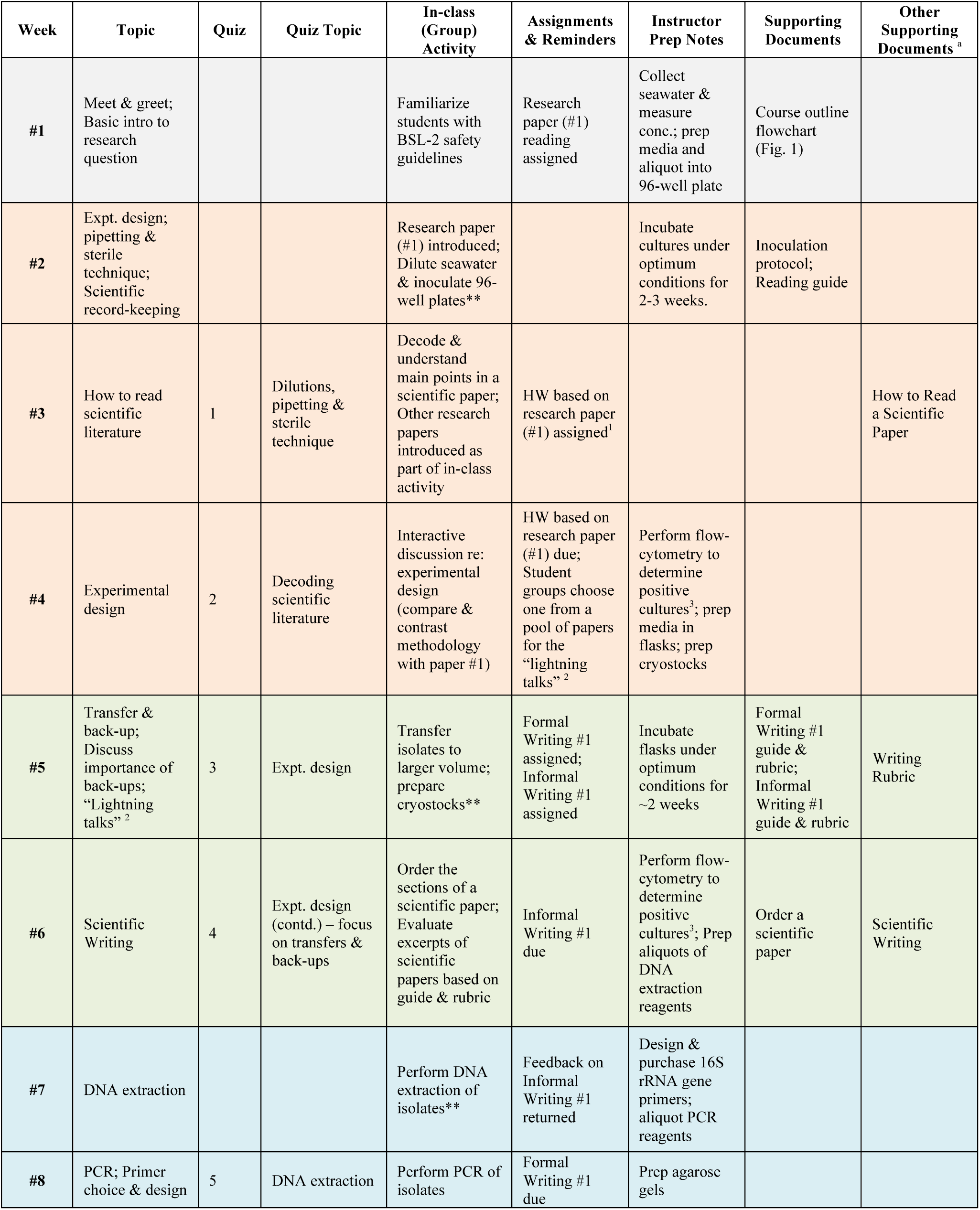
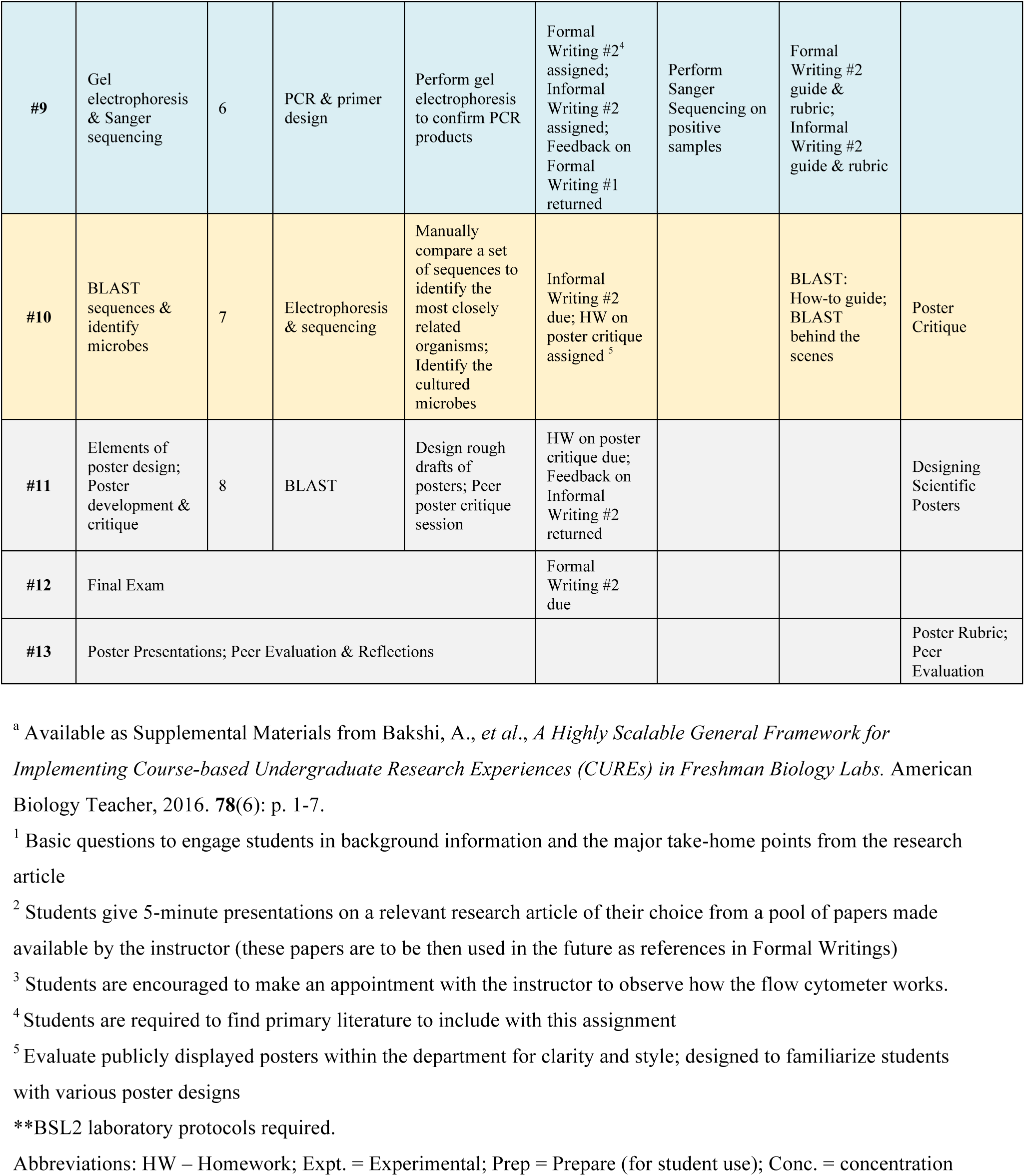
The mCURE framework. Activities, associated assessments, faculty instructions, and the relevant supporting documents are detailed week-by-week. The various segments of the course are color coded (grey, orange, green, blue, and yellow), consistently with the flowchart in Figure 1.

### Prerequisite student knowledge

Students are required to have basic prerequisite training and proficiency in biosafety level 1 (BSL1) organisms and safety practices (23). No other prerequisites are required, however, high school biology and chemistry are recommended. Students receive training in many of the basic biology skills that they will utilize in other contexts and receive training in biosafety level 2 (BSL2) protocols (see Safety Issues, below).

### Learning Outcomes

In addition to the learning objectives outlined below, the format of the mCURE sections incorporate aspects of three high-impact practices: Undergraduate Research, Collaborative Assignments, and Writing Intensive (24).

By the end of the semester, students should be able to:

1. Properly handle and isolate microorganisms using serial dilutions with the HTC protocol;
2. Extract DNA and amplify 16S rRNA genes from pure cultures;
3. Use databases such as BLAST to identify unknown microorganisms;
4. Describe the relationship between the research objectives, the HTC approach, and the experimental design;
5. Read and interpret relevant articles from the primary literature;
6. Communicate the methods, results, and implications of their research to both scientific and non-scientific audiences.

## PROCEDURE

A summary of the basic approach for the mCURE is shown in **Figure 1**, along with a week-by-week breakdown of activities, materials, and prep notes in **Table 1**.

### Materials

The required equipment and chemicals have been previously published (22). Briefly, because most highly abundant aquatic microorganisms have oligotrophic lifestyles, occur in low cell densities (< 10^7^ cells/mL), are very small (< 1μm), and will not grow on solid media, the cultivation approach makes use of liquid media, and cell growth is measured using a benchtop flow cytometer (e.g., the Millipore Guava easyCyte). The primary marine medium recipe, MWH1, and our flow cytometer settings, are provided in **Appendices 1** and **2**, respectively.

Alternative media recipes and preparation instructions are available elsewhere (14, 18). To avoid trace-metal contamination, all reusable cultivation vessels are made of polycarbonate plastic and acid-washed in 10% HCl. Other major items include a thermocycler and PCR reagents, electrophoresis equipment and a gel viewing system (e.g., Bio-Rad Gel Doc), a DNA quantification system (e.g. Qubit, ThermoFisher), DNA extraction kits (MoBio PowerWater), pipettes/tips, and incubators. The only differences in the established protocol equipment (22) for the mCURE sections are the requirement for a biosafety cabinet and disposable 2.1 mL 96-well plates (Thermo Nunc A/S). For those without access to some or most of this equipment, we provide alternatives in the Discussion.

### Student instructions

#### Segment 1 (*orange* in Figure 1, Table 1)

During the first two weeks of class, students are introduced to the overall mCURE approach, pipetting, and trained in BSL2 safety protocols. Each group of two students then dilutes their sample and inoculates seven wells of a 96-well plate (**Appendix 3**) containing the medium. An eighth well is inoculated with sterile media as a contamination control. Thus, a 24-student section initiates culturing in a 96-well plate. The plate is incubated at *in situ* temperature (based on time/place of sampling) for 2-3 weeks and then checked for growth using flow cytometry. During the incubation weeks, student assignments focus on introducing effective reading of scientific literature and on the experimental design and its rationale (**Table 1**).

#### Segment 2 (*green*)

Each group selects 1-2 positive cultures (wells with > 10^4^ cells/mL) for transfer into larger volume growth flasks and creates cryostocks for culture preservation in 10% DMSO (**Appendix 4**). In our experience, most groups usually have at least one positive well to transfer. Those groups with no growth in any of their wells select an unused positive well from another group. Inoculated flasks are incubated for two weeks at the same temperature as before. During the interim, students are introduced to scientific writing and give “lightning talks” (**Table 1**).

#### Segment 3 (*blue*)

Groups select at least one flask that shows growth and extract DNA (**Appendix 5**). In the three mCURE semesters detailed here, the majority of groups in any given section observed growth in at least one flask. Groups with no growth in any of their flasks use part of another group’s culture for extraction. Note that this introduces redundancy in the final identification results. Over the next two weeks, students amplify the 16S rRNA genes from their extracted DNA using PCR (**Appendix 6**) and confirm the amplification product with gel electrophoresis. Successful amplicons are then sequenced (possibly off-campus, e.g., the RTSF Genomics Core at Michigan State University).

#### Segment 4 (*yellow*)

Students learn to assemble forward and reverse sequence reads into a contig and identify their isolate using the NCBI BLASTN portal (**Appendix 7**). Briefly, reads from both the forward and reverse primer, as well as the overlapping contig (if any), are searched against the GenBank nt database with and without the exclusion of uncultured/environmental samples. The % identity, Query coverage, E-value, and GenBank # for the top five BLAST hits are recorded for all searches and isolates. Interpretation and contextualization of the results, including the similarity of isolates generated by the students to those in the database, occurs via discussion with knowledgeable faculty/teaching assistants. These results become part of their final poster presentation.

### Faculty instructions

#### Segment 1 (*grey*, *orange* in **Figure 1, Table 1**)

Prior to the beginning of the course, instructors must prepare the following:

1) Collect seawater (≥ 1L) and measure the concentration of bacterioplankton using flow cytometry (**Appendix 2**). The students use this initial concentration to calculate the dilution factor required to inoculate ~1-5 cells per well. Collection should occur as proximately to inoculation as possible to avoid microbial community change via bottle effects.
2) Prepare the low-nutrient media (**Appendix 1**; ~200 mL per plate; 1 plate/12 groups). Aliquot ~1.7 mL of media into each well of the 96-well plate just before class and allow time for equilibration to incubation temperature.
3) Select ~12-15 scientific articles (examples in **Appendix 8**) relevant to the project and create a reading guide for one of them for class discussion (sample: **Appendix 9** for (12)). The students may select one of the remaining papers for their lightning talks (**Table 1**, weeks #4-5), and use them as references for their formal writing assignments.

Because of the incubation period (2-3 weeks) for the initial inoculations, we recommend that Segment 1 involve at least one “holiday week” (**Table S1**). At the end of the incubation period, instructors count cells in the 96-well plate and record the well numbers positive for growth. Since isolates will be unknown at this time, transfers from incubation plates to counting plates (22) should be completed in a biosafety cabinet.

#### Segment 2 (*green*)

Prior to the start of this segment, instructors must prepare more medium, aliquot 50 mL into 125 mL flasks, and prepare cryotubes with DMSO. Prepare as many flasks and cryotubes as the number of wells that show growth (with some extra on hand in case of spillage). Students should have access to a biosafety cabinet in which to handle all cultures. At the end of the 2-week incubation, instructors count flasks to determine growth and record cell concentrations for student use. For the scientific writing discussion, we have made an activity (**Appendix 10**) that familiarizes students with the content in various sections of a paper (12).

#### Segment 3 (*blue*)

We recommend that instructors aliquot the required amount of DNA extraction reagents (**Appendix 5**-Power Water DNA Isolation Kit; Mo Bio Laboratories) and PCR reagents (**Appendix 6**-Taq, MgCl2, and buffer-ThermoFisher; 10mM AMRESCO dNTPs-VWR Life Sciences; 27F/1492R primers) for each group to prevent cross-contamination. For gel electrophoresis, gels are made with 1.5% agarose in DI MilliQ-filtered water. We suggest making an appropriate amount of agarose in a flask for each section, and allowing it to solidify until class time. Then, prior to the start of class, the instructor can melt the agarose in the flask and have it ready for students to pour their own gels. We recommend gels contain enough wells that each student has 1-2 wells to practice loading sample dye before loading their PCR product into one of the remaining wells. Students combine 1 μL loading dye with 5 μL PCR products for imaging. We typically employ a Lambda or 1 kb ladder. Gels are stained with SYBR green (1x) and imaged using the Bio-Rad Gel Doc.

#### Segment 4 (*yellow*)

Before the BLAST lab, instructors need to have all successful 16S rRNA gene amplicons sequenced from a facility of their choice using both forward and reverse primers (we use 27F and 1492R, but this can be specified by the instructor-see (25) for additional options); the resulting sequences should be made available where the students can access them. Label each sequence with the sample number and whether it is a “forward” or a “reverse” read. We recommend the “BLAST behind the scenes” activity (**Appendix 11**) to introduce students to the concept of sequence analysis. We have included the relevant lecture materials on molecular characterization (**Appendix 12**) to aid the instructor. Briefly, we introduce PCR, the importance of primers in PCR, describe the presence of conserved sequences flanking the hypervariable regions within 16S rRNA genes, and how the primers must be designed to recognize the conserved portion of the rRNA genes and amplify the hypervariable region they flank. We then discuss how Sanger sequencing can be used to read the DNA code and compared to other previously sequenced organisms using BLAST.

Finally, instructors need to prepare for a poster session at the end of the semester, including organizing space for poster boards, display tables, and printing facilities. However, for grading purposes, we recommend that the student groups present their posters electronically in class. During this time, other students and the instructor can offer constructive criticism for the students to incorporate into the final printed version of the poster.

Based on our experience implementing this mCURE for several semesters, we anticipate at least 1-2 protocol failures per semester; hence, flexibility is built into the framework (**Tables 1**, **S1**). Despite our anticipation of some failures and correcting these in subsequent semesters (e.g., students failing to properly transfer and freeze their samples), each new semester has presented us with new and different failures (e.g., flow cytometer reagents on back-order, failed PCRs due to old reagents). Many non-experimental activities, such as the lightning talks, can be easily inserted at different points in the course, amended to take less time, or even completely eliminated. Similarly, other related activities may be added, such as peer review of initial formal writing drafts and using social media for science outreach (*e.g.* we use the Twitter and Instagram hashtag #LSUCURE for all CURE efforts in the Department of Biological Sciences at LSU; **Table S1**). If feasible, we recommend adding the following enhancements to further engage students in the course: (i) taking students on a field trip, such as a one-day research cruise to collect water samples; (ii) demonstrating the use of “behind the scenes” equipment, such as the flow cytometer, capillary sequencer, and/or modern microscopes used to image bacteria.

### Suggestions for determining student learning

The mCURE is an authentic research experience and therefore one important component is communication of student findings for both scientific and non-scientific communities. Thus, assessment of student learning is largely split between the students successfully completing the protocols and the final poster presentation (**Table 2**). In order to complete the entire project, students need to be able culture bacterioplankton with the HTC protocol, passage cultures to larger volumes, extract DNA from these cultures, then successfully amplify and identify 16S rRNA gene sequences. The final poster and presentation requires students to state the aims of the project within the larger context of what is currently known about bacterioplankton in marine environments, outline the basic methodologies used, clearly present their results, and discuss these results in the context of their research question. Finally, the students suggest the next logical question to explore. Each of the laboratory and communication elements has multiple forms of evaluation (**Table 2** and **Appendices**).

**Table 2.** Determination of Student Learning.

| Learning Outcome (artifact) | Assessment Method(s)* |
| --- | --- |
| 1. Properly handle and isolate microorganisms using serial dilutions with the HTC protocol (isolated organisms). | Informal Writing 1 (Appendix 14), Formal Writing 1 (Appendix 16), successful completion of the protocols, results presented in the final poster (Appendix 21). |
| 2. Extract DNA and amplify 16S rRNA genes from pure cultures (16S rRNA gene amplicons). | Informal Writing 2 (Appendix 15), Formal Writing 2 (Appendix 17), successful completion of the protocols, results presented in the final poster (Appendix 21). |
| 3. Use databases such as BLAST to identify unknown microorganisms (taxonomic identity). | Formal Writing 2 (Appendix 17), successful completion of the protocols, results presented in the final poster (Appendix 21). |
| 4. Describe the relationship between the research objectives, the HTC approach, and the experimental design. | Formal Writing 2 (Appendix 17), final poster (Appendix 21). |
| 5. Read and interpret relevant articles from the primary literature. | Lighting Talks (Appendix 19), Formal Writing 2 (Appendices 17), final poster (Appendix 21). |
| 6. Communicate the methods, results, and implications of their research to both scientific and non-scientific audiences (poster). | Lighting Talks (Appendix 19), final poster (Appendix 21). |

\*Rubrics for both the writing assignments have been published previously (10).

### Sample data

Fall 2015, spring 2016, and fall 2016 average cultivability (13) was 9.9, 2.8, and 12%, respectively. These cultivability numbers generally match the success rate of other HTC experiments (14) and demonstrate a significant improvement over “traditional” methods (11). The number of unique pure cultures that survived successive transfers and were positively identified at the end of each course was 28 (fall 2015), 13 (spring 2016), and 23 (fall 2016). In total, mCURE sections isolated 43 unique bacterioplankton during the first three semesters reported herein. Some courses isolated taxa identified in a previous mCURE, so the overall total was smaller than the sum of the individual semesters. Many of the isolates have close relationships to organisms previously cultured using HTC in the Thrash lab and other labs, as indicated by taxonomic affiliations to strains with “LSUCC”, “HTCC”, “HIMB”, or “IMCC” designations (**Table 3**). Importantly, many isolates represent abundant marine clades (14), thus the results validate the mCURE approach to produce valuable cultures with similar efficacy as HTC experiments conducted under more typical laboratory settings. Additional results are provided in **Appendix 13**.

**Table 3.** Bacteria cultured by mCURE students

| **Closest unique cultured relative** | **Major Taxonomic Group** |
| --- | --- |
| **Arthrobacter sp. 210_2** | Actinomycetales; Actinobacteria |
| **Marinobacterium sp. IMCC1424** | Actinomycetales; Actinobacteria |
| **Microbacterium esteraromaticum strain V45.13** | Actinomycetales; Actinobacteria |
| **Microbacterium sp. Ni17** | Actinomycetales; Actinobacteria |
| **Nocardioides exalbidus strain DS1-2B** | Actinomycetales; Actinobacteria |
| **Nocardioides hwasunensis strain XH199** | Actinomycetales; Actinobacteria |
| **Alteromonadales bacterium 3tb13** | Alteromonadales; Gammaproteobacteria |
| **Alteromonas macleodii** | Alteromonadales; Gammaproteobacteria |
| **Alteromonas tagae** | Alteromonadales; Gammaproteobacteria |
| **Marinomonas sp. SS8** | Alteromonadales; Gammaproteobacteria |
| **Porticoccus hydrocarbonoclasticus** | Alteromonadales; Gammaproteobacteria |
| **Pseudoalteromonas phenolica** | Alteromonadales; Gammaproteobacteria |
| **Pseudoalteromonas sp. A-3** | Alteromonadales; Gammaproteobacteria |
| **Shewanella sp. 49WBP** | Alteromonadales; Gammaproteobacteria |
| **Bacillus sp. L1(2012)** | Bacillales; Firmicutes |
| **Burkholderiales bacterium LSUCC0118** | Burkholderiales; Betaproteobacteria |
| **Limnobacter sp. MYOU6** | Burkholderiales; Betaproteobacteria |
| **Haliaceae bacterium LSUCC0247** | Haliaceae; Gammaproteobacteria |
| **Gamma proteobacterium SF293** | OM182; Gammaproteobacteria |
| **Gamma proteobacterium IMCC15037** | OM252; Gammaproteobacteria |
| **Gammaproteobacteria bacterium LSUCC0258** | OM252; Gammaproteobacteria |
| **Gammaproteobacteria bacterium LSUCC0272** | OM252; Gammaproteobacteria |
| **Marine gamma proteobacterium HTCC2080** | OM60/NOR5; Gammaproteobacteria |
| **Agrobacterium sp. TSH97** | Rhizobiales; Alphaproteobacteria |
| **Anderseniella baltica** | Rhizobiales; Alphaproteobacteria |
| **Anderseniella baltica strain BA141** | Rhizobiales; Alphaproteobacteria |
| **Rhizobium sp. MSSRF QS100** | Rhizobiales; Alphaproteobacteria |
| **Bacterium HIMB11** | Rhodobacterales; Alphaproteobacteria |
| **Rhodobacteraceae bacterium LSUCC0246** | Rhodobacterales; Alphaproteobacteria |
| **Rhodobacteraceae bacterium LSUCC0259** | Rhodobacterales; Alphaproteobacteria |
| **Roseobacter sp. strain WM2** | Rhodobacterales; Alphaproteobacteria |
| **Altererythrobacter ishigakiensi** | Sphingomonadales; Alphaproteobacteria |
| **Erythrobacteraceae bacterium LSUCC0210** | Sphingomonadales; Alphaproteobacteria |
| **Erythrobacteraceae bacterium LSUCC0236** | Sphingomonadales; Alphaproteobacteria |
| **Erythrobacteraceae bacterium LSUCC0240** | Sphingomonadales; Alphaproteobacteria |
| **Erythrobacteraceae bacterium LSUCC0267** | Sphingomonadales; Alphaproteobacteria |
| **Bacterium MH1** | Vibrionales; Gammaproteobacteria |
| **Vibrio chagasii** | Vibrionales; Gammaproteobacteria |
| **Vibrio pelagius** | Vibrionales; Gammaproteobacteria |
| **Vibrio proteolyticus** | Vibrionales; Gammaproteobacteria |
| **Vibrio sp. 0208F3** | Vibrionales; Gammaproteobacteria |
| **Vibrio sp. PaH3.31d** | Vibrionales; Gammaproteobacteria |
| **Vibrio sp. TP187** | Vibrionales; Gammaproteobacteria |

### Safety issues

Since the curriculum involves isolating unknown organisms, students must be proficient in BSL1 safety techniques prior to taking the course. All activities that involve handling live microorganisms should occur under BSL2 safety protocols, as outlined by the JMBE Biosafety Guidelines for Handling Microorganisms in the Teaching Laboratory (23). The specific activities requiring BSL2 protocols are indicated in Table 1. Additional safety measures must be taken for faculty during washing and preparation of medium mixture bottles and growth flasks. See (22) for more details.

## DISCUSSION

### Field testing

Here we report results from mCURE sections offered during the fall 2015 and 2016 semesters in Biology 1207 (Honors: Biology Laboratory for Science Majors) and spring 2016 in Biology 1208 (Biology Laboratory for Science Majors I). There were four sections per semester taught by two graduate teaching assistants (two sections each), with up to 28 students per section. Biology 1207 is only offered in the fall semester and consists of a total of four sections. Multiple (12-50) sections of Biology 1208 are offered every semester, a few of which are typically offered as CUREs as outlined in our previous publication (10); students do not know when they register for this course if their section will be in a CURE or traditional format. We note that these previous sections of the mCURE were conducted with a BSL1 safety protocol. The current protocol offered in this manuscript has been updated with BSL2 safety measures in response to recommendations by ASM (23). In each of these sections, some fraction of student groups (pairs) were capable of successfully implementing the protocols from start to finish, while others had failures that required they use cultures, DNA, or PCR products from other groups. In general, we found that roughly a third of the groups could successfully complete the entire workflow (however, failure at any given step did not preclude students from progressing to the next step, albeit with successful cultures from a different group). This represents only one of the learning outcomes. Other learning outcomes (**Table 2**) could be achieved regardless of students experiencing failure at different stages (detailed below).

### Evidence of student learning

We provide evidence of student learning with example summative assessment of grade distributions (**Fig. 2**), physical data (PCR products-**Fig. 3**), qualitative results of successfully completed bacterioplankton isolation (**Table 3**), and examples of the range of student communication outcomes (**Table 4**, **Appendix 22**).

**Figure 2.**
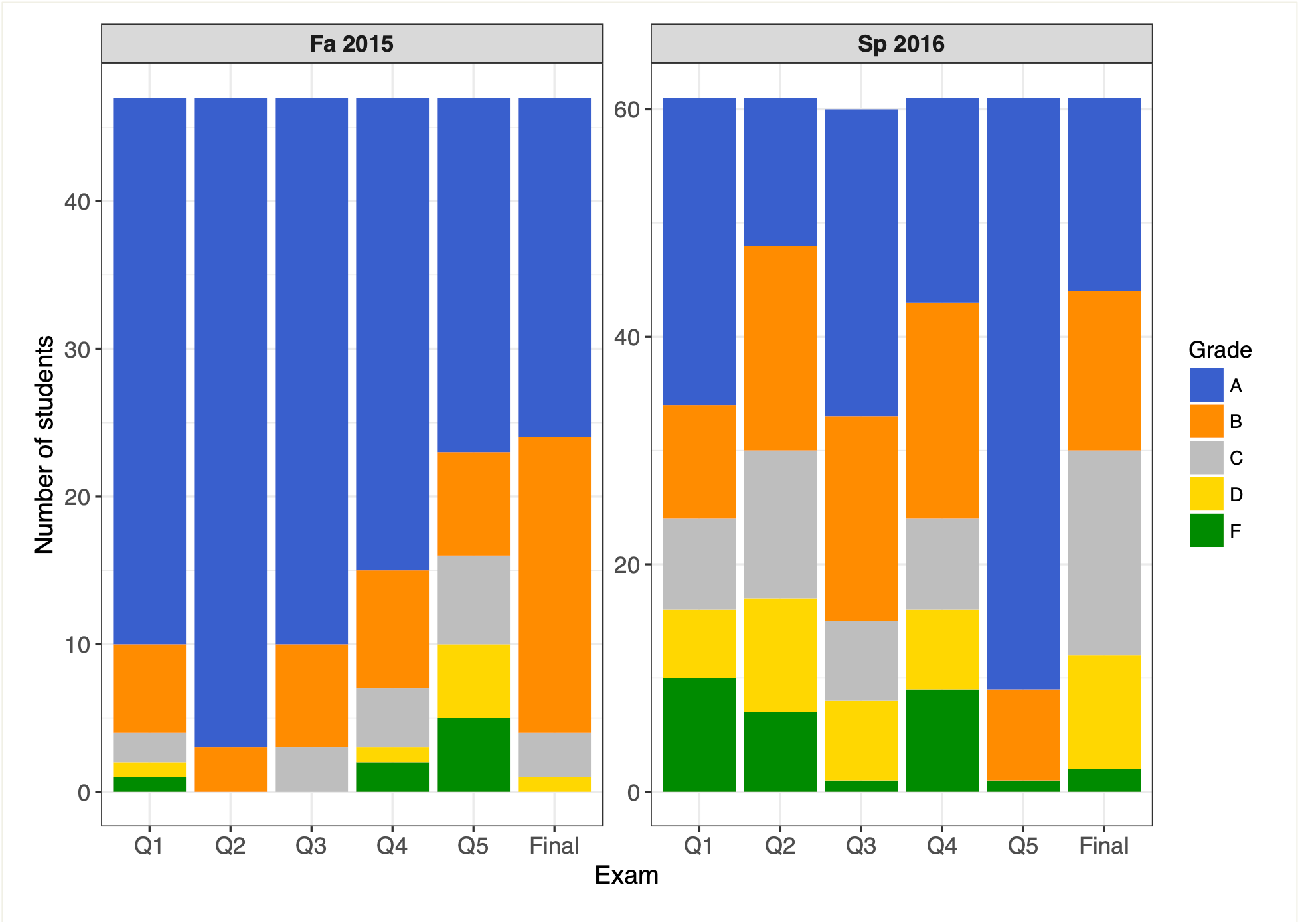
Grade distributions for two sections of mCURE students during each of two semesters in the 2015-2016 school year. **Fall 2015** consisted of ~50 Honors College students majoring in biology. The topics for the five quizzes (Q1-Q5) were as follows: Q1 = Safety, Controls; Q2 = Experimental design, Scientific writing; Q3 = DNA extraction; Q4 = PCR; Q5 = Gel electrophoresis, Purpose of sequencing, Primer design. **Spring 2016** consisted of ~60 mostly non-biology major students. The topics for the five quizzes (Q1-Q5) were as follows: Q1 = Dilutions, Pipetting, Safety, Controls, Scientific writing; Q2 = Experimental design, Dilution, Pipetting, Controls; Q3 = DNA extraction; Q4 = PCR, Primer selection/design, Gel electrophoresis; Q5 = Purpose of sequencing, Sequence analysis. The grades for both semesters were assigned based on the following score criteria: A = 90-100%; B = 80-90%; C = 70-80%; D = 60-70%; F = <60%.

**Figure 3.**
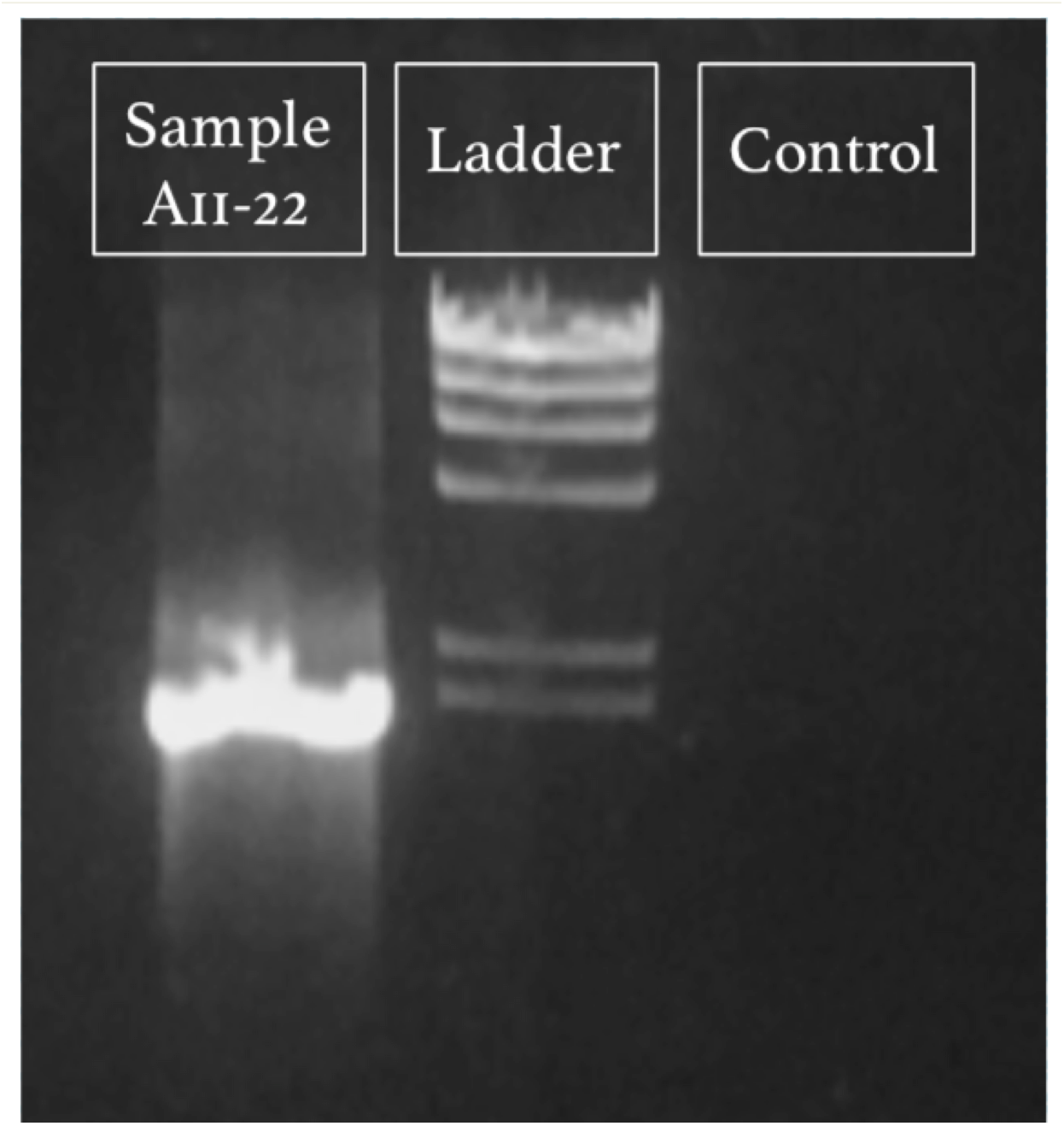
Example gel electrophoresis image of a successful 16S rRNA gene PCR amplification from fall 2015. Lanes labeled according to contents: “Sample A11-22” is the amplicon from isolate DNA (expected size 1466 bp); “Ladder” is Lambda HindIII digest ladder (NEB N3012S), with the lowest visible band at 2027 bp; “Control” is the negative control (water).

**Table 4.** Excerpts from students’ posters describing the bacteria they cultured. Students were expected to identify and describe major points of interest regarding the bacteria they cultured, supported by scientific literature references, relate that information back to the experimental design, and identify a future direction for their work. Minor spelling and grammatical errors have been fixed when reformatting the excerpts to fit the format of this table.

|  | **Excerpts about the cultured organisms from students' posters** |
| --- | --- |
| **Excellent** | *Pseudoalteromonas phenolica* was originally found in 2003 by Alim Isnansetyo and Yuto Kamei in the waters near the islands of Japan. Species in the genus *Pseudoalteromonas* are typically heterotrophic but [some may] be oligotrophic, which is what our experiment is designed to culture. . . . The most significant attribute of this organism, though, is that it produces anti-methicillin-resistant *Staphylococcus aureus* (MRSA) substances (Isnansetyo and Kamei 2003). . . . Because this species, *Pseudoalteromonas phenolica*, produces anti-MRSA substances, more focus should be put on how effective these substances are against *Staphylococcus aureus*. Experiments should be done to see if this species can be grown easily in large quantities to be produce [the] antibiotic.<br>*Several interesting attributes of the cultured bacteria and major points of significance are explained in detail with proper citations; future directions identified, and information related back to the experiment students conducted; demonstrates thorough understanding of experimental design.* |
| **Good/**<br>**Acceptable** | *Pseudoalteromonas phenolica*, found from B5-1, is significant because it can be used to treat MRSA, a bacterium that can cause skin infections, infected wounds and even pneumonia, that has resistance to many known antibiotics. It could possibly be used in a pharmaceutical product to treat illnesses caused by MRSA in the future. [In the future, we could] use the cryostocks to culture the organism . . . to confirm its identity . . . and attempt to find if our strain has anti-MRSA properties.<br>*Organism's important attribute of scientific interest identified and its significance described but not cited; future directions identified, and information related back to the experiment students conducted; demonstrates thorough understanding of the experimental design.* |
| **Needs**<br>**Improvement** | [*Pseudoalteromonas phenolica*] was first cultured in a lab near Tokyo, Japan in 2003. Strains are currently being researched for their antibiotic properties on anti-methicillin-resistant *Staphylococcus aureus*. 4 out of 11 groups at LSU cultured a *P. phenolica* showing that it is abundant in the Gulf of Mexico and readily grows through HTC. [Future directions include] identifying biological markers, studying its contributions to the ecosystem, and finding industrial, medical, and pharmaceutical applications.<br>*Organism briefly described and important attributes mentioned without* |
|  | *expanding upon their significance or proper citations; future directions identified, but information not related back to the experiment conducted; demonstrates incorrect understanding of the experiment conducted (several students that semester characterized <u>P. phenolica</u> because not many cultures were initially successful, thus a few groups had to share the same initial broth cultures for the molecular analysis steps).* |

**Figure 2** details the grade distributions across two sections from each semester during the 2015-2016 school year, composed of students with differing levels of academic preparation. The fall 2015 sections consisted of Honors College students majoring in biology, many of whom were already familiar with basic laboratory techniques. These students did not perform the original dilution of the seawater before inoculation. This class generally performed well on quizzes, which tested their proficiency in one or two of the major topics covered in the prior week of the course. Nearly the entire class received a grade of either A or B on the cumulative final exam (**Appendix 18**, **Fig. 2**). In spring 2016, we offered the mCURE in BIOL 1208R. Spring is the “off” semester for this course such that students enrolled in it usually are not biology majors or experienced some barrier to their enrollment or completion of the course in the preceding fall semester. This semester we asked students to perform their own seawater dilution. Many students found this difficult, as reflected in the Q1 and Q2 scores (**Fig. 2**). However, we note that by the final exam most students were proficient in these calculations. At the end of the semester, ~75% of the class received a passing grade (A-C) on the final exam, which is typical for the traditional lab sections during the spring semester of this course.

In addition to demonstrating their knowledge on summative assessments, students became proficient in laboratory techniques (Learning Outcomes 1-2) as evidenced by the vast majority of student groups in both semesters who successfully extracted DNA from cultures and performed PCR (e.g., **Fig. 3**). By the end of the semester, students were expected to understand and interpret primary literature related to their research and describe their cultured microbe in the final poster. Thus, the posters partially address Learning Outcomes 3-6, with other writing assignments providing additional training (**Table 2**). **Table 4** provides excerpts from student posters describing their isolated organism. The top performing students included detailed description of scientific literature related to their organism and proposed future experiments to expand our knowledge about their isolate. Their writing was concise while including all important and relevant details and showed a thorough understanding of the experimental design. We provide examples of formal writing assignment 2, lightning talks, and student posters in **Appendix 22** (shared with permission from the students).

### Possible modifications

We appreciate that many instructors may wish to implement the mCURE design but may not have access to some of the more expensive equipment used in our protocol. Here are a few modifications to circumvent some of these restrictions. Instructors can replace flow cytometry with direct microscopic counts, e.g., as in some of the earlier iterations of the HTC protocol (12). For those without access to either a flow cytometer or a fluorescence microscope, the protocol can still be completed using traditional agar-plate based methods. Our media can be prepared with agar (22) or replaced with a classic marine medium like Difco 2216 (BD). Although solid media generally select for different taxa than liquid media, for the purposes of a basic biology laboratory, this may not matter. After streaking a seawater sample on plates, individual colonies can be picked, grown up in liquid culture to increase cellular mass, or directly processed through DNA extraction. Colony PCR (26) may also be an attractive alternative identification method, particularly because this also eliminates the time and cost associated with DNA extraction. These last two steps may also help adapt the overall protocol for shorter time frames, e.g., academic quarters instead of semesters. Please note that our protocol uses low-nutrient and low-carbon media that typically selects for non-pathogenic, oligotrophic marine bacterioplankton (14). The use of rich media and plate-based methods may increase the risk of cultivating pathogenic organisms. Finally, for those interested in freshwater environments, the same protocol can be conducted with freshwater media, either artificial (18, 27) or natural (28).

## SUPPLEMENTAL MATERIALS

**Table S1.** Example of a real implementation schedule of the idealized template in Table 1.

**Appendix 1.** MWH1 marine medium recipe

**Appendix 2.** Flow cytometry parameters

**Appendix 3.** Inoculation Protocol

**Appendix 4.** Transfer and Cryostock Protocol

**Appendix 5.** DNA Extraction Protocol

**Appendix 6.** PCR protocol

**Appendix 7.** BLAST How-to Guide

**Appendix 8.** Suggested literature

**Appendix 9.** Reading Guide

**Appendix 10.** Ordering a Scientific Paper

**Appendix 11.** BLAST Behind the Scenes

**Appendix 12.** Molecular biology lectures (as PowerPoint slides). If readers would like assistance in developing other PowerPoints such as these, please contact the lead author.

**Appendix 13.** Supplemental results

**Appendix 14.** Informal Writing 1

**Appendix 15.** Informal Writing 2

**Appendix 16.** Formal Writing 1

**Appendix 17.** Formal Writing 2

**Appendix 18.** Sample Final Exam

**Appendix 19.** Lightning talk rubric

**Appendix 20.** Quizzes

**Appendix 21.** Poster rubric

**Appendix 22.** Example Student Assignments

Appendix 23. Example lightning talk and instructions (as PowerPoint slides).

## ACKNOWLEDGMENTS

We wish to thank Dr. Chris Gregg, Ann Dickey-Jolissaint, and Brooke Trabona for coordinating the labs and prepping many of the lab materials behind the scenes. David Morris and Celeste Lanclos served as additional graduate and undergraduate TAs, respectively. Andrew Flick and Dr. Paige Jarreau were instrumental in incorporating social media assignments into the curriculum. The Socolofsky Microscopy Center provided images of some of the microbes cultured by students and gave students tours of the facility. We thank the crew of the *R/V Acadiana* and Murt Conover at the Louisiana Universities Marine Consortium for assisting with logistics and field training. We are grateful to Dean Cynthia Peterson for her support of CUREs in the Department of Biological Sciences. Funding for this project was awarded through the Student Excellence Fee from the College of Science.

